# T492I mutation alters SARS-CoV-2 properties via modulating viral non-structural proteins

**DOI:** 10.1101/2023.01.15.524090

**Authors:** Xiaoyuan Lin, Zhou Sha, Jakob Trimpert, Dusan Kunec, Chen Jiang, Yan Xiong, BinBin Xu, Zhenglin Zhu, Weiwei Xue, Haibo Wu

## Abstract

The historically dominant SARS-CoV-2 Delta variants and the currently dominant Omicron variants carry a T492I substitution within the non-structural protein 4 (NSP4). Based on a combination of *in silico* analyses, we predicted that the T492I mutation increases the transmissibility and adaptability of the virus. We confirmed this hypothesis by performing competition experiments in hamsters and in human airway tissue culture models. Furthermore, we show that the T492I mutation also increases the replication capacity and infectiveness of the virus, and improves its ability to evade antibody neutralization induced by previous variants. Mechanistically, the T492I mutation increases cleavage efficiency of the viral main protease NSP5 by enhancing enzyme-substrate binding, resulting in increased production of nearly all non-structural proteins processed by NSP5. Importantly, T492I mutation suppresses the viral RNA associated chemokines in monocytic macrophages, which may contribute to the attenuated pathogenicity of Omicron variants. Our results highlight the importance of the NSP4 mutation in the evolutionary dynamics of SARS-CoV-2 and identify a novel target for the development of broad-spectrum antiviral agents.

## INTRODUCTION

Severe acute respiratory syndrome coronavirus 2 (SARS-CoV-2) has triggered a global public health crisis since 2019 [1,2]. The virus has a moderate mutation rate [3–5], but is in a rapid evolution due to the large number of human infections. Genomic variations and the birth of adaptive mutations enabled the rapid spread of several distinct variants of concern (VOCs) [6–9]. The S mutation N501Y [10–13] and the nucleocapsid (N) mutation R203K/G204R [14] play an essential role in the increased infectivity and transmissibility of VOC Alpha, a historical predominant lineage in the first half of 2021. Another S mutation, L452R, was demonstrated to increase infectivity, transmissibility and resistance to neutralization of VOC Delta, which replaced Alpha as the dominant lineage in the second half of 2021 [15,16]. The present ongoing VOC Omicron contains a strikingly high number of mutations [17], especially in the S protein. These spike adaptations in Omicron allow SARS-CoV-2 variants to dominate the current pandemic [18]. Functional analyses of these important mutations provide clues to the rising of VOCs and the evolution of SARS-CoV-2, enrich the understanding of molecular and cellular mechanisms of SARS-CoV-2 infection, and guide the development of effective intervention strategies.

Previous work mainly focused on SARS-CoV-2 spike mutations, because the S protein mediates the receptor recognition and fusion processes and is the main target for the development of vaccines and neutralizing antibodies [19,20]. Nonstructural proteins (NSPs) have been studied far less than S protein, although NSPs account for more than 70% of the SARS-CoV-2 genome. NSPs constitute the viral replication and transcription complex (RTC) [21], interact with host proteins during the early coronavirus replication cycle, and initiate the biogenesis of replication organelles [22–24]. In addition, NSPs have been reported to be associated with coronavirus pathogenicity [25–27], and mutations on NSPs may lead to attenuated virulence of Omicron compared with previous lineages [28]. Recently, Chen et al. reported that ancestral SARS-CoV-2 isolate carrying the Omicron S gene caused similar severe disease in mice [29], suggesting that non-Spike Omicron mutations may be primarily responsible for the reduced pathogenicity. Of the NSPs, NSP4 is involved in the formation of double-membrane vesicles (DMVs) closely associated with viral replication [30] and, together with NSP3 and NSP6, redirects host inner membranes into replicating organelles [31,32]. Meanwhile, NSP4 induces extensive mitochondrial structural changes, the formation of outer membrane macropores, and the release of mitochondrial DNA-loaded inner membrane vesicles [33]. Furthermore, NSP4 is known as a strong predictor of mortality in patients with severe COVID-19 [34,35]. The functional importance of NSP4 suggests that accumulative mutations in this protein may contribute significantly to the phenotypic changes of SARS-CoV-2 along its evolutionary history.

In this study, we focused on the NSP4 mutation T492I, which is shared by the Delta dominant sub-variant (21J) and all Omicron sub-variants. We tracked the evolution of T492I based on all documented SARS-CoV-2 genomic sequences. The results showed that the incidence frequency (IF) of T492I has increased since January 2021 and spread rapidly with the emergence of Delta 21J lineage and all Omicron lineages. The mutation has fixed in the worldwide pandemic. Comprehensive evolutionary analysis revealed evidences of positive selection supporting the transmission advantage and adaptiveness of T492I. Through experimental studies conducted in cell lines, hamsters, and human airway tissue models, we identified and validated increased infectivity and fitness of the 492I variant. Further, we found a decreased sensitivity of virus carrying the 492I mutation towards the sera of hamsters infected with wild-type SARS-CoV-2 virus, suggesting an enhanced immune evasion capability. This finding is supported by the fact that 492I virus showed increased fitness relative to T492 virus after the initiation of the global vaccination program. Moreover, we observed a decrease in the severity of 492I virus infected hamster lung tissues, which has been confirmed by epidemic surveys and global clinical data statistics. In summary, our results suggest that NSP4 T492I substitution plays an essential role in the rapid spread and enhanced immune evasion capability of SARS-CoV-2 VOCs, from Delta to Omicron.

## RESULTS

### Evolutionary trajectory of NSP4 mutation T492I

Delta is the dominant VOC in the second half of 2021, and Omicron is the dominant VOC from early 2022 to present (Figure 1A, B). Comparison of the genomic sequences of VOCs showed that the mutation T492I (C10029T) in the nonstructural protein 4 (NSP4) is shared by Delta and Omicron. Another two VOCs Lambda and Mu also bear T492I (Figure 1A), based on the analysis of the sequences and the information provided by NextStrain [36] and GISAID [37]. These suggest a potential importance of this mutation to the fitness and transmission of SARS-CoV-2. We evaluated the evolutionary trajectory of this mutation to assess its contribution to the spread of Delta and Omicron. The phylogenetic tree of variants and the statistics of the incidence frequency (IF) showed that T492I used to appear in early variants (19B and 20B, in the beginning of 2021) and had an increase in IF before the emergence of Delta (Figure 1A, B). Among the three Delta lineages, 21A, 21I and 21J, the lineage 21J carried T492I, but 21A and 21I did not (Figure 1A). The Delta 21J lineage had a higher transmission frequency than 21A and 21I, and it was the predominant strain when Delta reached a nearly fixed frequency (Figure S1A, B). Mutation T492I took a percentage of nearly 100% in Omicron variants (Figure S1C, D), which replaced Delta as the predominant VOC starting in 2022 (Figure S1E, F). To date, T492I is a fixed NSP4 mutation all over the world (Figure 1A).

**Figure 1.**
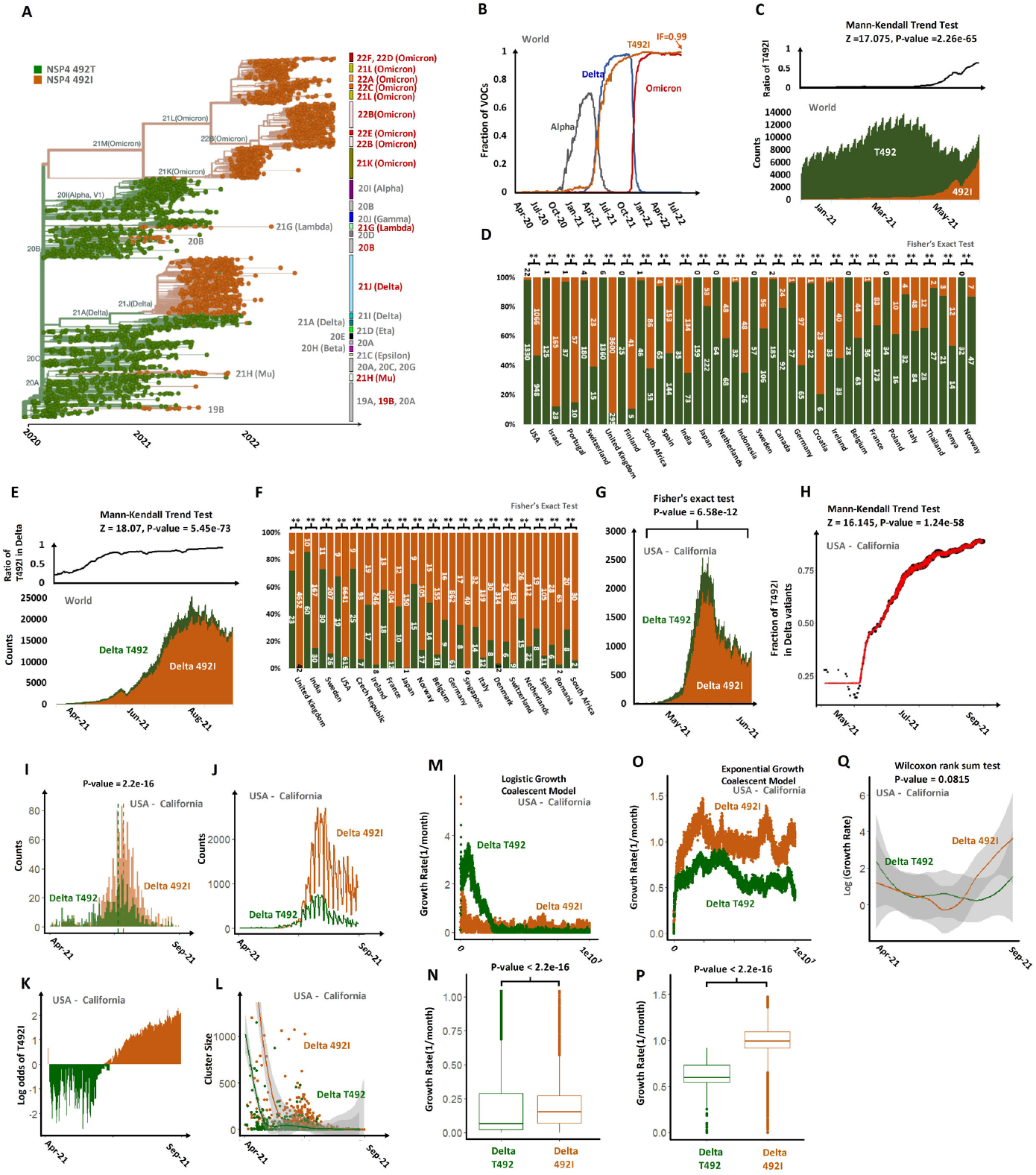
Evidences supporting the adaptiveness of NSP4 mutation T492I. (A) A phylogenetic tree of SARS-CoV-2 strains showing the rapid spread of T492I and associated VOCs, marked in red. The phylogenetic tree was acquired from NextStrain and curated. (B) IF changes of T492I and three VOCs (Alpha, Delta and Omicron) up to November 2022. (C) The upper panel shows the changes in the ratio of the T492I with the result of the Mann-Kendall trend test listed at the top. The lower panel shows the weekly running counts of the T492 and 492I variants from January 2021 to June 2021. (D) Comparison of the fractions of the T492 and 492I variants at two time points separated by a gap of more than 2 weeks in different countries, corresponding to Table S1. ‘**’ denotes significance in Fisher’s exact test. The counts of identified samples are marked within bars. (E) Comparison between Delta T492 and Delta 492I variants. Legend in (E) follows (C). (F) Comparison of the fractions of the Delta T492 and Delta 492I variants at two time points separated by a gap of more than 2 weeks in different countries, corresponding to Table S2. (G) Comparison of weekly running counts between T492 Delta variants and 492I Delta variants in a regional subdivision, USA-California. The result of Fisher’s exact test at the time interval bracketed is shown at the top. (H) The fitted trend of the change in the fractions of the T492I in the Delta variants in USA-California, with the result of the Mann-Kendall trend test listed at the top. The red folded line is the maximum likelihood estimate with a non-decreasing constraint. The dot size reflects the number of identified samples on that day. (I-L) Temporal distribution of T492 Delta and 492I Delta phylogenetic clusters in the California State of USA. (I) Counts of California T492 Delta and 492I Delta clusters when first detected and over time. (J) Numbers of T492 Delta or 492I Delta samples collected over time. (K) Log odds of the frequency of 492I/T492 in Delta variant over time. (L) Relationship between cluster size (Y axis) and the date when the first sample was collected within a cluster (X axis). (M-Q) Comparison of phylodynamic growth rates between T492 Delta and 492I Delta variants collect in USA-California from April 2021 to September 2021. (M) and (N) are the comparison of the growth rates across states in a logistic growth coalescent model. (O) and (P) are those in an exponential growth coalescent model. (M) and (O) are scatter diagrams (X axis denote the state) and (N) and (P) are box diagrams. (Q) is a comparison of growth rates (logged in the plot) over time simulated in the skygrowth coalescent model.

### T492I confers increased transmission fitness

For an evaluation of the hypothesis that T492I confers increased transmission fitness, we compared IF between the T492 and 492I variants over a six-month interval from the temporal vicinity of the initial rise, from January 2021 to June 2021. The comparison was performed in multiple geographically restricted contexts. The results showed that T492I had a global increase in IF (Figure 1C, S1G, Table S1). The increase was significant in Fisher’s exact test of the fractions of pairs of lineages on the onset and beyond the day two weeks later, in Mann-Kendall trend test and in isotonic regression analysis. There are 24 countries with a statistical significance in the three statistical tests and 23 of them have an increased IF (Binomial Test, P-value 2.98e-06, Figure 1D, S1H, I, Table S1). Sixty-eight regional subdivisions were with a significance in the three statistical tests, 53 of which had increased IF (Binomial Test, P-value 4.116e-06, Figure S1J, K, Table S1). To avoid the effect of other mutations, we used the same method to compare IF between T492 and 492I in Delta variants (21J vs 21A + 21I) in a six-month interval from the initial rise of Delta (April 2021 to September 2021). We still found that the Delta T492I variants had a significant IF increase in three geographical scales, including the global scale (Figure 1E, S1L), the country level and the regional subdivision level. There are 19 countries with an increased IF out of 19 countries with a statistical significance in the three statistical tests (Binomial Test, P-value 3.815e-06, Figure 1F, S1M, N, Table S2) and 55 regional subdivisions with an increased IF out of 56 regional subdivisions with a statistical significance in the three statistical tests (Binomial Test, P-value 1.582e-15, Table S2).

The strains collected in State of California, USA showed a significant increase in IF of T492 in the three statistical tests referred above (Figure 1G, H). For a further evaluation of the adaptiveness of T492I, we compared the phylogenetic clusters of Delta T492 variants and Delta 492I variants collected in this regional subdivision in a six-month interval as referred above (April 2021 to September 2021). Clusters of Delta 492I were first detected later than Delta T492 clusters (Figure 1I, P-value = 2.2e-16). There were nearly equal number of Delta 492I clusters (178) and Delta T492 clusters (173), but the former were 191% larger than the latter on average (P-value = 0.004, Figure 1J). The sampling frequency of T492I mutation continuously increased in Delta variants (Figure 1K), and the earliest detected clusters were larger than those detected later (Figure 1L). Simulation with a logistic coalescent model and an exponential growth coalescent model both showed that the growth rates of 492I Delta variants were higher than those of T492 Delta variants (Figure 1M-P). The estimated mutational selection coefficient (s) of T492I was 1.376 for the logistic growth coalescent model and 0.669 for the exponential growth coalescent model, assuming that the T492 Delta variants grew exponentially at a rate of r and the Delta 492I variants grew exponentially at a rate of r(1+s). The Delta 492I variants also showed a higher growth rate than the Delta T492 variants with weak significance in a simulation with the skygrowth coalescent model (Figure 1Q). These suggest the adaptiveness in the transmission of the T492I mutation.

### T492I endows SARS-CoV-2 with higher competitiveness and infectivity

To validate the impact of NSP4 T492I mutation on SARS-CoV-2, we constructed a 492I variant based on the USA_WA1/2020 SARS-CoV-2 sequence (GenBank accession No. MT020880), and conducted competition experiments using a Syrian hamster model as previously described [14,38,39]. The results showed that a higher 492I to T492 ratio was observed at 3 and 5 days post infection (dpi), indicating a sustained advantage of 492I virus over T492 virus in hamsters (Figure 2A, S2A). Similar competition experiments were conducted in a human airway tissue culture model. We found that, when airway tissues were infected with control and 492I virus at a ratio of 1:1, the ratio of 492I variant to T492 virus increased from 1 to 5 dpi (Figure 2B). After infection of airway tissues with control virus and 492I variant in a 3:1 or 9:1 ratio, 492I variant rapidly overcame the initial deficit and showed an advantage over the control virus (Figure 2C, D). These data indicate that 492I variant could rapidly outcompete the T492 virus in human airway tissues.

**Figure 2.**
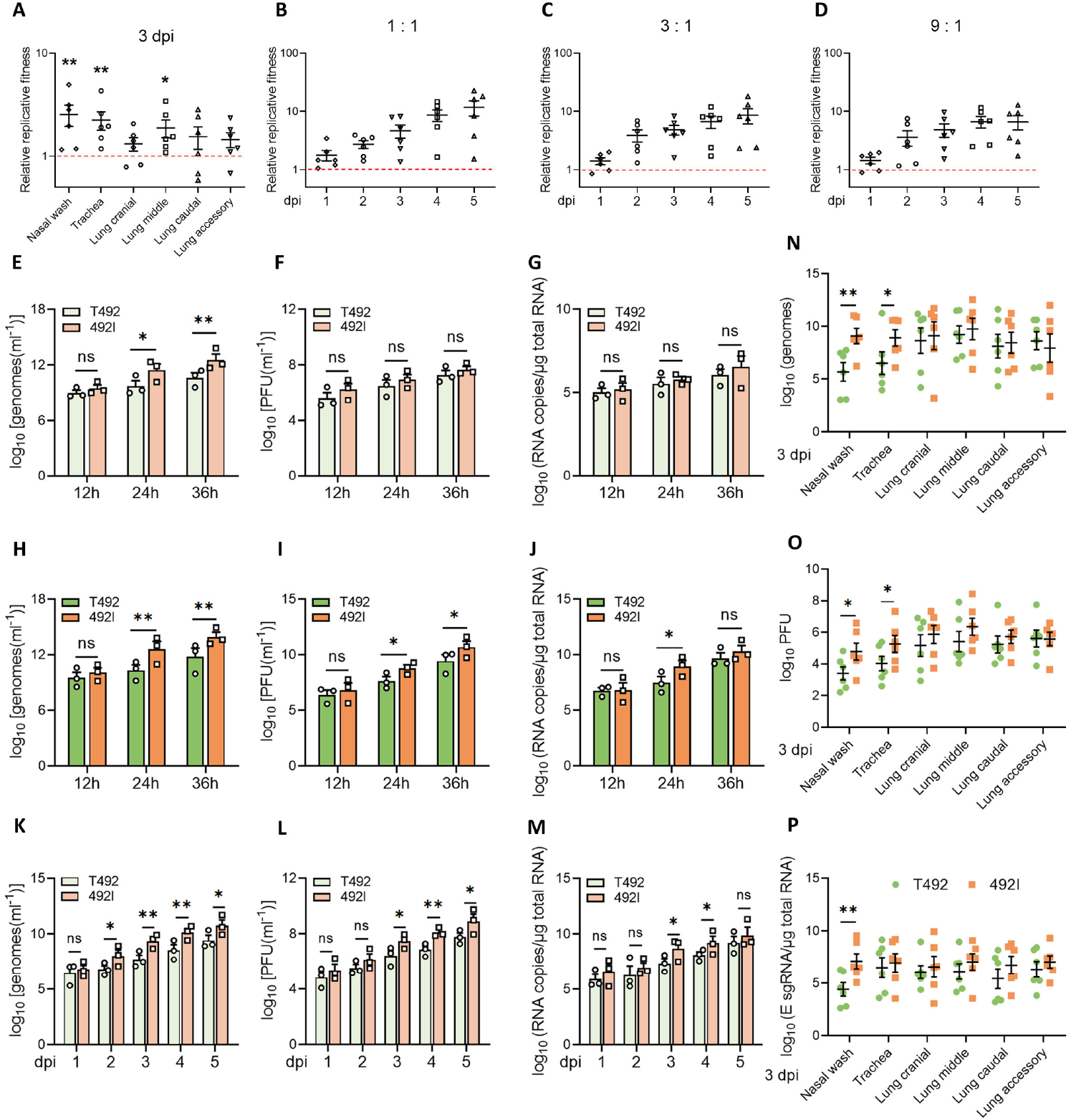
SARS-CoV-2 bearing a T492I mutation has higher competitiveness and infectivity. (A) Hamsters were infected with T492 and 492I virus at a mixture of 1:1. The relative amounts of T492 and 492I viral RNA in nasal wash, trachea and lung samples were detected by RT-PCR and Sanger sequencing at 3 dpi. Log10 scale was used for the Y-axis. Dots represent individual hamsters (n = 6). (B-D) Mixture of T492 and 492I virus was inoculated into human airway tissue cultures at an initial ratio of 1:1 (B), 1:3 (C) or 1:9 (D) with a MOI of 5. Relative amounts of T492 and 492I viral RNA were detected by qRT-PCR and Sanger sequencing for 5 consecutive days. Log10 scale was used for the Y-axis. Dots represent individual tissue culture (n = 6). (E-J) Genomic RNA levels (E, H), PFU titres (F, I), and E sgRNA loads (G, J) of T492 and 492I viruses were evaluated in Vero E6 (E-G) and Calu-3 (H-JF) cell lines. Vero E6 or Calu-3 cells were infected with T492 or 492I virus at a MOI of 0.01. Viral replication, infectious titres and subgenomic RNA were detected by plaque assay and qRT-PCR, respectively. Experiments were performed in triplicate. (K-M) T492 or 492I virus was inoculated into human airway tissue cultures at a MOI of 5. Genomic RNA levels (E, H), PFU titres (F, I), and E sgRNA loads (G, J) were detected for 5 consecutive days. Experiments were performed in triplicate. (N-P) Hamsters were infected with 2×10^4^ PFU of T492 or 492I virus. Genomic RNA levels (N), PFU titres (O), and E sgRNA loads (P) in nasal wash, trachea and lung samples were detected at 3 dpi. Dots represent individual hamsters (n = 6). All data are presented as the mean ± s.e.m.. *, p<0.05, **, p<0.01. Abbreviation: ns, nonsignificant.

Next, we tested the replication and infectivity of 492I variant in Vero E6 monkey kidney cell line and Calu-3 human lung epithelial cell line. The results showed that 492I variant replicated with higher extracellular viral RNA than T492 virus at 24 and 36 hours post infection (hpi) (Figure 2E). The infectivity of virus was measured by PFU titres and viral subgenomic RNA (E sgRNA) loads. As a result, there were no significant differences in PFU titres and E sgRNA loads between 492I variant and T492 virus in Vero E6 cells (Figure 2F, G); however, in Calu-3 cells, 492I virus produced significantly higher extracellular viral RNA than the control (Figure 2H), and similar trends in infectious titres and E sgRNA loads were observed (Figure 2I, J). In line with these results, we found that extracellular viral RNA, PFU titres and viral E sgRNA loads of 492I variant were significantly higher than those of T492 virus in human airway tissue cultures (Figure 2K-M). Furthermore, replication and infectivity tests were conducted using a hamster model. As a result, although hamsters infected with different viruses showed similar weight loss during the infection (Figure S2B), viral RNA (Figure 2N, S2C), infectious titres (Figure 2O, S2D), and E sgRNA loads (Figure 2P, S2E) in nasal wash and trachea samples from hamsters infected with 492I variant at 3 and 5 dpi were significantly higher than in controls. Taken together, these results indicate that T492I mutation enhances viral replication and infectivity of SARS-CoV-2.

### T492I mutation increases the cleavage efficiency of NSP5

We explored strategies by which the T492I mutation affected viral properties. SARS-CoV-2 gene order is similar to that of other known coronaviruses, with the first two open reading frames (Orf1a and Orf1b) encoding all non-structural proteins [21]. NSP4 protein is a 500-amino acid protein with four predicted transmembrane domains, and T492I mutation is located next to the NSP4|5 cleavage site (also known as the N-terminal autoprocessing site of NSP5) (Figure 3A). NSP5 (M pro or 3CL-pro) is thought to cleave the viral polyprotein at multiple distinct sites, yielding nonstructural proteins NSP5–NSP16 that support viral replication and disable host defense [40,41]. It has been reported that the amino sequence of the N-terminal autoprocessing site is essential for the cleavage efficiency of NSP5. Inhibition of the NSP5-mediated cleavage prevents NSPs production and results in impaired viral replication [21,42,43]. Based on these, we speculated that T492I mutation might affect the cleavage efficiency of NSP5. To address this hypothesis, we constructed a fusion protein substrate with a linker sequence based on the NSP4|5 cleavage site in-between a 3×Flag-His tag and the full length GFP (Figure 3B). In this gel-based cleavage assay, we found that the cleavage efficiency of NSP5 on T492I linker substrate was higher than that on control linker substrate, and that cleavage was observed as early as 30 min in T492I group (Figure 3C). These data imply that T492I mutation enhances the cleavage efficiency of NSP5.

**Figure 3.**
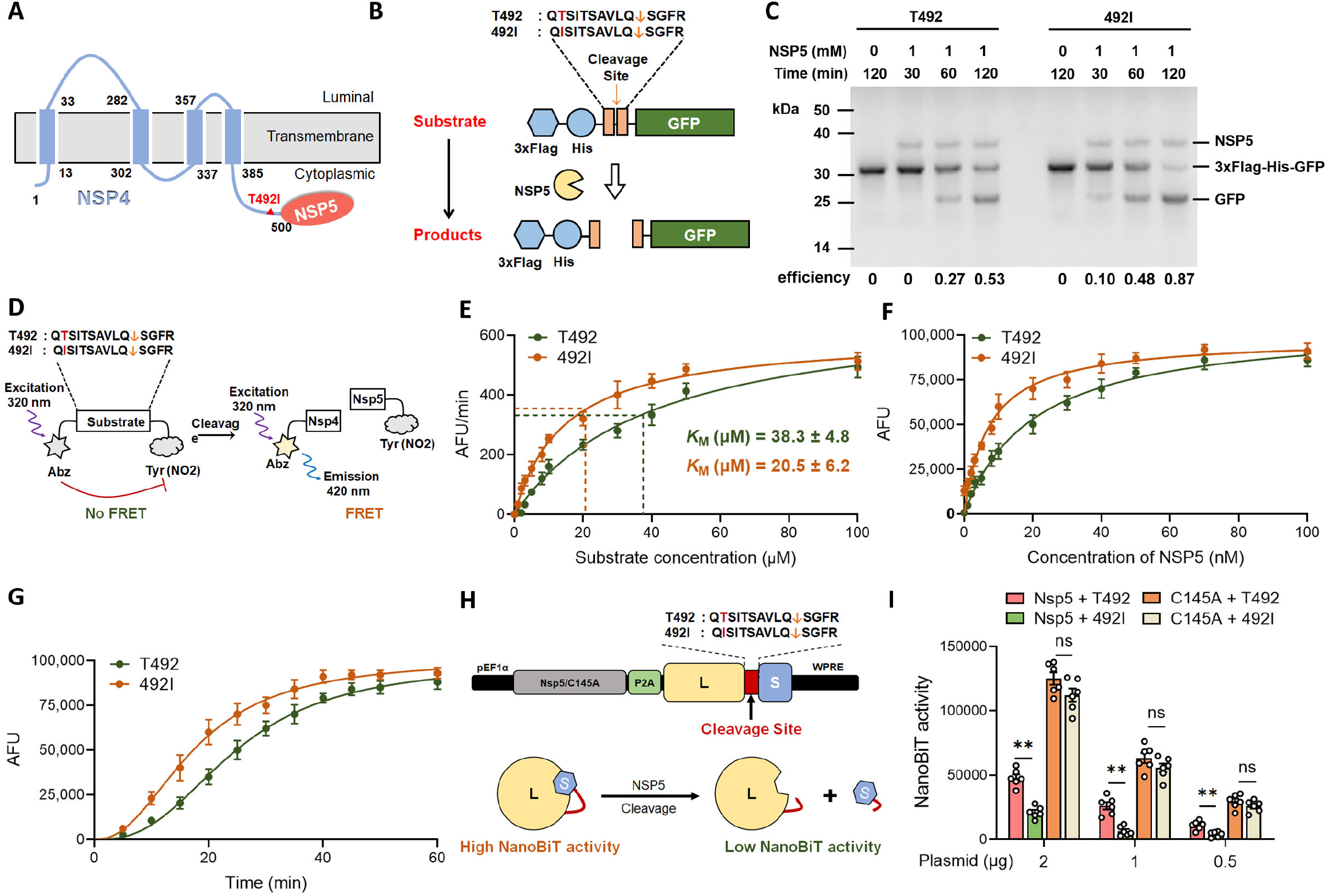
T492I mutation increases the cleavage efficiency of NSP5. (A) Predicted structure of SARS-CoV-2 NSP4, a 500-amino acid protein with four predicted transmembrane domains. The mutation site (at residue 492) is indicated with a red triangle. (B) Schematic diagram of the gel-based cleavage assay. (C) Gel-based assay for substrate cleavage at 1mM NSP5 over time is shown. Cleavage of the substrate yields two products: 3×Flag-His (~4 kDa, not visible on the gel) and GFP (~27 kDa). The molecular weight of the uncleaved substrate is approximate 31 kDa and NSP5 is approximate 35 kDa. (D) Schematic diagram of the FRET-based protease assay. The fluorescence of Abz in the uncleaved substrate is quenched by N-tyrosine. If NSP5 cleaves the substrate, the fluorescent signal increases accordingly. (E-G) Comparison of the enzyme kinetics of NSP5 protease on T492 and 492I substrate. KM values were determined by measuring the initial reaction rates over a range of concentrations and then plotting against the substrate concentration (E). The value of fluorescence over the increase of NSP5 concentrations is shown (F). Fluorescence intensities were detected at 20 min after the start of reaction. The value of fluorescence at 10 μM NSP5 concentration over time is shown (G). Data are presented as the mean ± s.e.m.. Experiments were performed in triplicate. (H) Schematic diagram of NanoBiT assay. NanoBiT is a luciferase complementation reporter comprised of a Large BiT (L) and a Small BiT (S). The L and S fragments are connected by a linker containing the T492 or 492I cleavage site. Cleavage of the linker results in a low luciferase activity. (I) HEK-293T cells were transfected with plasmid expressing NSP5/C145A and NanoBiT reporter as indicated, and cell lysates were collected for luciferase assay at 36 h post transfection. Experiments were performed in triplicate. Data are presented as the mean ± s.e.m.. **, p<0.01. Abbreviation: ns, nonsignificant.

To compare the enzyme activity of NSP5 on different substrates, we performed fluorescence resonance energy transfer (FRET)-based assays using a synthesized 14 amino acid-long peptide substrates based on the NSP4|5 cleavage site. The FRET peptide contains a fluorophore (2-aminobenzoyl, Abz) at its N-terminus and a quencher (N-Tyrosine) at its C-terminus. NSP5 cleaves the substrate and releases fluorophore from the proximity of quencher, resulting in an increase in fluorescent signal (Figure 3D). Using the FRET peptides, we determined that the Michaelis constant (KM) of NSP5 was 38.3 ± 4.8 μM for control substrate, and 20.5 ± 6.2 μM for 492I substrate (Figure 3E). NSP5 cleavage of 492I substrate was more efficient than cleavage of control substrate (2.37-fold increase in kcat/KM; Table S3). The enzyme activity data of increase in fluorescence over increasing NSP5 concentrations (Figure 3F), as well as the data of increase in fluorescence over time at 10 μM NSP5 concentration (Figure 3G), indicated that T492I mutation increased the cleavage efficiency of NSP5 on the NSP4|5 cleavage site.

Further, we optimized a NanoBiT assay to validate our findings. NanoBiT is comprised of a Large BiT (L) and a Small BiT (S), which can complement each other to form a functional luciferase protein. We connected L and S fragments with a linker containing the NSP4|5 cleavage site. NSP5 cleaves the linker and causes L and S fragments to dissociate, resulting in a low luciferase activity (Figure 3H). The result showed that the NanoBiT activity collected from NSP5 cleavage of 492I linker group was significantly lower than that of control linker group (Figure 3I). Moreover, the catalytically inactive form of NSP5 (C145A, Cys145 was converted to Ala), which was unable to process the NSP4|5 cleavage site, was introduced to the system. We found that NSP5-C145A mutant showed no significant difference in the cleavage of control and 492I linkers (Figure 3I). These data confirm that T492I mutation of substrate facilitates NSP5 cleavage efficiency, and the enzymatic activity is remains dependent on the C145 catalytic-site.

### T492I mutation enhances the NSP5-substrate contact

Next, we asked whether T492I mutation affected enzyme-substrate contact. Using a proximity ligation assay, we showed that 492I substrate can approach to the NSP5 enzyme more effectively than T492 substrate (Figure 4A). Results collected from *in vitro* co-immunoprecipitation assays showed that NSP5 enzyme bound to Flag-tagged T492 and 492I substrates in a dose-dependent manner. In line with previous results, the binding affinity of NSP5 to the 492I substrate was stronger than that to the control substrate (Figure 4B).

**Figure 4.**
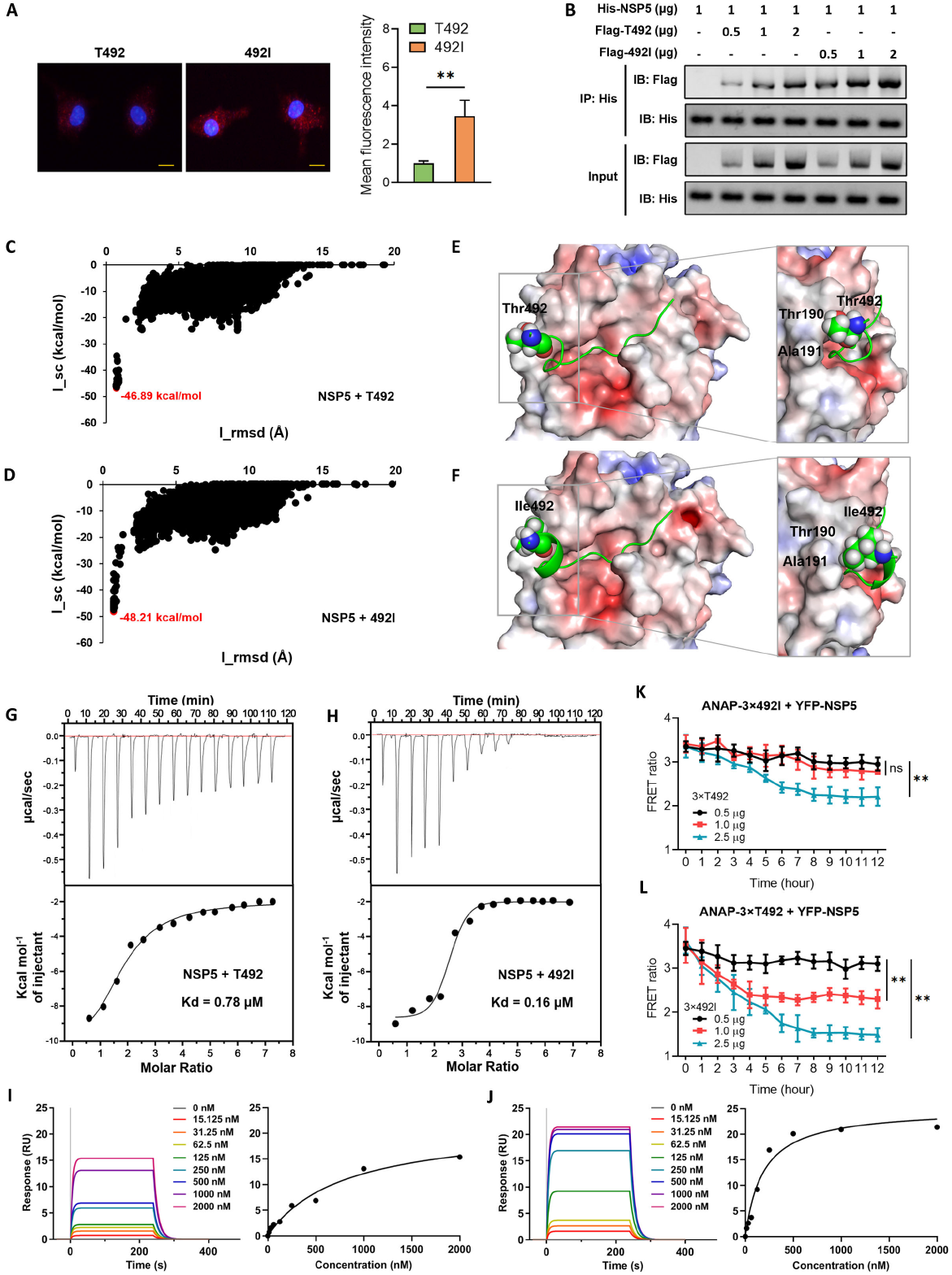
T492I mutation enhances the NSP5-substrate contact. (A) Proximity ligation assay of NSP5 and T492I substrates. Vero E6 cells were transfected with plasmids encoding Flag-tagged T492 or 492I substrates. Twelve hours later, cells were infected with SARS-CoV-2 virus at a MOI of 0.01 for another12 hours. Cells were fixed and the fluorescence intensities were detected using anti-Flag and anti-NSP5 antibodies. The histogram on the right panel represents the mean value of fluorescence intensities. Scale bar = 10 μm. (B) Co-immunoprecipitation assay of NSP5 and Flag-tagged T492I substrates. His-NSP5 together with Flag-T492I or Flag-492I plasmids were co-transfected to HEK-293T cells as indicated. After 24 hours, cell lysates were harvested and immunoprecipitated using an anti-Flag antibody. The lower panel represents the immunoblot analysis of whole cell lysates. (C, D) Computational modeling for the binding of T492 and 492I substrates to SARS-CoV-2 NSP5. The Rosetta docking funnels of the T492 (C) and 492I (D) substrate to NSP5. The red plots have the lowest docking interface score (I_sc) with the interface root-mean-square deviation (I_rmsd) ≤ 4 Å. (E, F) Predicted binding modes of T492 (E) and 492I (F) substrates to NSP5 corresponding to the lowest docking score in (C) and (D). The cartoon (green) and surface (APBS-generated electrostatic) models were used for the substrate and NSP5, respectively. The residues Thr492 and Ile492 were shown in sphere model. (G, H) ITC-binding curve between the T492 (G) and 492I (H) substrate with NSP5. (I, J) Left panels are the representative SPR sensorgrams of the response over time when increasing concentrations of T492 (I) or 492I (J) substrates were injected over purified NSP5 protein immobilized on biosensor chip. Right panels are the kinetic analysis of T492 substrate-NSP5 (I) and 492I substrate-NSP5 (J) complexes. The Kd values were determined by the steady-state affinity of each concentration. (K, L) HEK-293T cells were transfected with CMV-3×492I(TAG)-NSP5-YFP (K) or CMV-3×T492(TAG)-NSP5-YFP (L) plasmid, and then treated with 50 μM ANAP. After 18 hours, cells were transfected with different dose (0.5, 1.0 and 2.5 μg, respectively) of 3×T492 (K) or 3×492I (L) plasmid. Graph displaying the FRET ratios (YFP/ANAP) recorded from single cell images. Schematic diagram of this intracellular FRET assay was shown in Figure S3. Experiments were performed in triplicate. Data are presented as the mean ± s.e.m.. **, p<0.01. Abbreviation: ns, nonsignificant.

To understand the structural basis of different cleavage efficiencies, we determined the 3D structures of NSP5 binding to T492 and 492I substrates. The presence of docking funnel in Figure 4C and 4D indicated that control and 492I substrates were successfully docked with SARS-CoV-2 NSP5. For each complex, the structure with the lowest I-sc and I-rmsd ≤ 4Å from the docking trajectory was selected as a near-native model of the substrate binding to NSP5. The lowest I-sc for the T492 and 492I substrates bound complexes were −46.89 kcal/mol (Figure 4C) and −48.21 kcal/mol (Figure 4D), respectively. Compared with the control substrate, 492I mutant exhibited a relatively better binding affinity to NSP5, which is consistent with the trend of our *in vivo* and *in vitro* experiments on protease binding activity. Furthermore, the molecular mechanism underlying the stronger binding affinity of 492I was clarified by comparing the binding patterns between the control and 492I substrates to NSP5. It was clearly shown that when the amino acid Thr at substrate 492 mutated to Ile, the latter was more suitable to form hydrophobic contacts with residues Thr190 and Ala191 in the NSP5 substrate binding pocket (Figure 4E, F).

Furthermore, we performed isothermal titration calorimetry (ITC) and surface plasmon resonance (SPR) to determine the accurate dissociation rate constants (Kd) of NSP5/T492 and NSP5/492I complexes. The results obtained from ITC (Figure 4G, H) and SPR (Figure 4I, J) showed that the affinity of 492I substrate to NSP5 was about 4.13~5.78-times that of T492 substrate to NSP5. Additionally, we used FRET to demonstrate that 492I substrate had a competitive advantage for NSP5 in human embryonic kidney (HEK) 293 cells. ANAP-modified 3×492I substrate and YFP-tagged NSP5 were constructed for the intracellular FRET system (Figure. S3A, B). FRET ratio showed that the binding of 492I substrate to NSP5 was only slightly weakened in competition with high dose of T492 substrate (Figure 4K). On the contrary, a small amount of 492I substrate was able to hijack NSP5 from the NSP5-T492 substrate complex (Figure 4L). These data collectively reveal that T492I mutation enhances the enzyme-substrate contact, thereby increasing the protease activity of NSP5.

### T492I mutation alters SARS-CoV-2 properties by affecting non-structural proteins

NSP5 is the main protease of SARS-CoV-2 genome and plays a major role in the cleavage of viral polyproteins. NSP5 cleaves polyproteins to generate individual functional proteins, such as RNA-dependent RNA polymerase (NSP12), helicase (NSP13), endoribonuclease (NSP15) and other indispensable cofactors. We then detected the amount of NSPs cleaved by NSP5 in Calu-3 cells infected with control and 492I virus, respectively. The ELISA results showed that the protein levels of NSPs in 492I group were significantly higher than that in the control group, except for NSP4 and NSP8 (Figure 5A). This data validated that T492I mutation increased the cleavage efficiency of NSP5. Among the NSPs affected by T492I mutation, NSP4 and NSP6 have been reported to form a replication-transcription complex (RTC) along with a few host factors. RTC is associated with modified host ER membranes that generate convoluted membranes (CMs) and double-membrane vesicles (DMVs) for viral genome replication and transcription. To further confirm the impact of T492I mutation on viral life cycle, membrane rearrangement caused by viral infection was investigated by transmission electron microscopy. Analysis of virus-infected Calu-3 cells showed that T492I variant induced more extensive CMs and DMVs than control group (Figure 5B). This data is consistent with our previous result that T492I mutation enhanced the viral replication of SARS-CoV-2 (Figure 2E, H, K, N).

**Figure 5.**
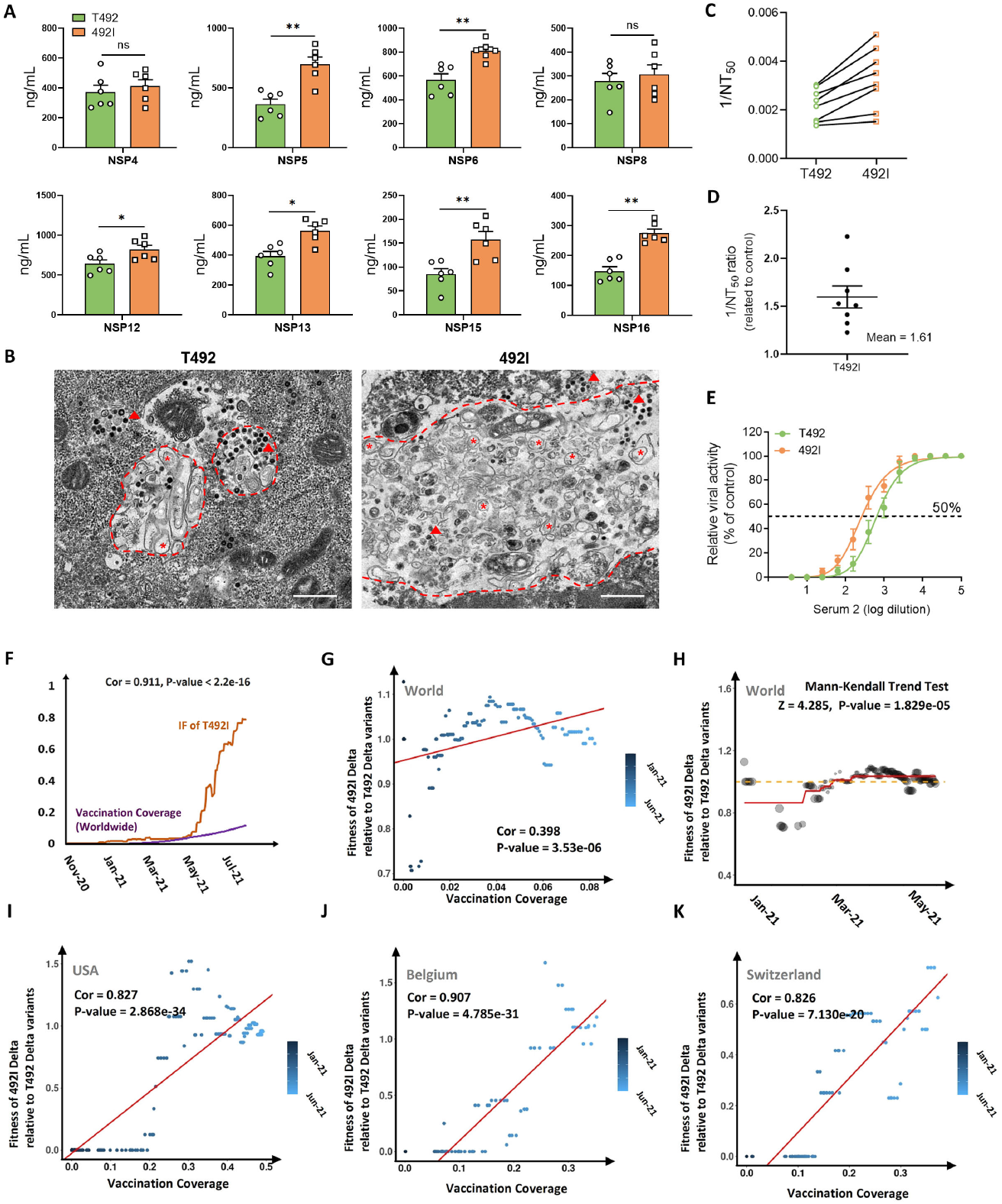
T492I mutation shows an association with strengthened resistance to neutralization and decreased disease severity. (A) Calu-3 cells were infected with T492 or 492I virus at a MOI of 0.01. After 48 hours, the cell lysates were harvested and subjected to ELISA analysis using specific NSP antibodies. (B) Calu-3 cells were infected with T492 or 492I virus at a MOI of 0.01. After 48 hours, cells were fixed and processed to transmission electron microscopy. Convoluted membranes are marked with red dotted lines, double-membrane vesicles are marked with red asterisks, and virus particles are marked with red arrowheads. Scale bar = 500 nm. (C) Neutralization assay of hamster sera against T492 and 492I virus containing a mNeonGreen reporter. 1/NT_50_ values were plotted. Dots represent sera from individual hamsters (n = 8). (D) 1/NT_50_ values of 492I virus to T492 virus are shown. Dots represent sera from individual hamsters (n = 8). (E) Neutralization curves of serum 2 from individual hamsters. The solid line represents the fitted curve, and the dotted line indicates 50% viral inhibition. (F-K) Statistical evidences that suggest an association between T492I and resistance to neutralization. (F) An image contemporarily showing the IF of T492I variants and the worldwide vaccination coverage. The correlation between them is 0.911 with statistical significance. (G, I-K) Correlation analysis results between the fitness of 492I Delta relative to T492 Delta variants (y-axis) and VC (x-axis) in the world (G), USA (I), Belgium (J) and Switzerland (K). The color of the dots corresponds to the date. (H) The fitted trend of the fitness of Delta 492I relative to Delta T492 variants in the world. The result of Mann-Kendall trend test are listed at the top. Data in (A, D, E) are represented as the mean ± s.e.m.. *, p<0.05, **, p<0.01. Abbreviation: ns, nonsignificant.

As previously reported, the endonuclease activity of NSP15 [44,45], and the methyltransferase activity of NSP16 [46–48], both serve to evade the innate immune response of host. We subsequently assessed the effect of T492I mutation on virus sensitivity to the neutralization serum. Neutralization titres of a panel of sera collected from hamsters infected with wild type virus were analyzed using the mNeonGreen reporter 492I virus, as previously reported [14]. In comparison with the control virus, neutralization titres of all sera against 492I virus were reduced by 1.22-to 2.38-fold (mean 1.61-fold) (Figure 5C, D). Serum 2 presented the lowest neutralization titres against 492I virus (Figure 5E). These data imply that T492I mutation of NSP4 endows SARS-CoV-2 variant with the ability to evade antibody neutralization.

Moreover, based on the statistics of the global vaccination data, we observed a significant correlation between the IF of T492I and the global vaccination coverage (VC) (Figure 5F), suggesting a possible correlation between T492I and immune evasion. Further, we evaluated the changes in fitness of Delta 492I relative to Delta T492 variants and the correlation between fitness and VC. Relative fitness was calculated based on the changes in the IF of Delta 492I and Delta T492 variants. The evaluation was performed in a global level and in different countries. We observed an increase of relative fitness of T492I and this increase was clearly correlated with an increase of VC not only in a global level (Figure 5G, H) but also in a country level (Figure 5I-K, Table S4). There were 23 countries with an increased relative fitness (Binomial Test, P-value 2.98e-06) and 22 countries with a positive correlation (Binomial Test, P-value 3.588e-05) out of 26 countries with a statistical significance both in Mann-Kendall trend test and correlation coefficient test (Table S4). These data support a strengthened resistance to neutralization of T492I mutation.

### T492I mutation shows an association with decreased disease severity

To understand the impact of T492I mutation on viral pathogenicity, we compared the lung lesions of hamsters infected with T492 or 492I virus. The result showed that hamsters infected with 492I virus exhibited less lung damage and pulmonary vascular congestion than the control group (Figure 6A). Interestingly, this reduction in pathogenicity was only shown at late stage of 5 dpi (Figure 6A, B). To validate this finding, we performed pulmonary function tests. We found that hamsters infected with T492 virus displayed increased functional residual capacity (Figure 6C), total lung capacity (Figure 6D), vital capacity (Figure 6E) and residual volume (Figure 6F) at 5 dpi than hamsters infected 492I virus. Importantly, hamsters infected T492 virus showed a lower forced expiratory volume in 100 ms (FEV_0.1_, Figure 6G), and possessed means of forced expiratory volume in 100 ms/forced vital capacity less than 0.49 (FEV_0.1_/FVC, equivalent to the FEV1/FVC index in human, a criterion routinely used for pulmonary function diagnosis, less than 0.49 means severe. Figure 6H). Collectively, these data suggest that 492I virus presents decreased lung pathologies and milder pulmonary function injury in hamsters compared to the control virus.

**Figure 6.**
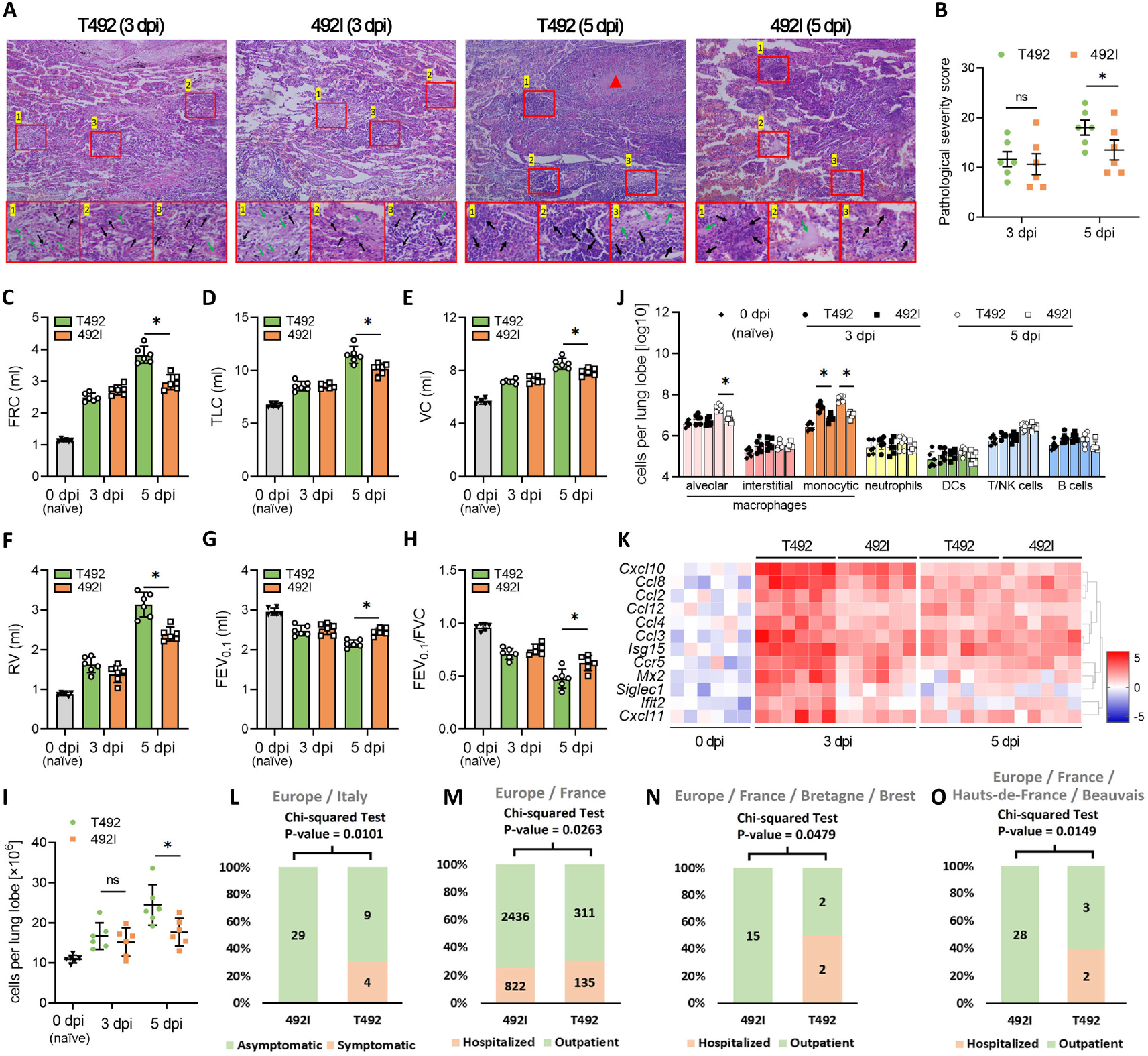
T492I mutation shows an association with decreased disease severity. (A) Haemotoxylin and eosin staining of lung sections collected at 3 and 5 dpi from hamsters infected with 2×10^4^PFU of T492 or 492I virus. The lower photographs are magnified images of the regions denoted by rectangles in the upper photographs. The lower panel shows bronchioles with aggregation of inflammatory cells (Black arrow) and surrounding alveolar wall infiltration (Green arrow). Red arrowheads indicate the alveolar parenchymal lesions. (B) Histopathology scoring of lung sections. Lung lobes were scored individually using the scoring system described in the methods. The scores of 5 slices from each hamster were added to determine the total pathology score per animal. (C-H) Pulmonary function tests of hamsters infected with T492 or 492I virus. Functional residual capacity (C), total lung capacity (D), vital capacity (E), residual volume (F), forced expiratory volume in 100 ms (G) and forced expiratory volume in 100 ms/forced vital capacity (H) were detected at 3 and 5 dpi. (I-J) Total leukocytes count (I) and specific cell clusters count (J) per lung lobe in hamsters infected with T492 or 492I virus and control group (naïve, 0 dpi). (K) Relative expression of monocytic macrophage-specific chemokines associated with viral RNA was detected by qRT-PCR. Coloration and point size indicate log2 fold change. (L-O) Prediction of the clinical outcomes of T492 Delta and 492I Delta variants. Collection sites are listed at the top of the figures. Y axis shows the ratios between lineages with opposite patient statuses. Lineage numbers are provided. The significance of changes in ratios were tested by chi-squared test. Dots represent individual hamsters (n = 6). Data in (B-J) are presented as the mean ± s.e.m.. *, p<0.05. Abbreviation: ns, nonsignificant.

To obtain the dynamics of pulmonary response during infection, total leukocytes recruitment (Figure 6I) and specific cell clusters (Figure 6J) were counted by flow cytometry as previously described [49]. Cell type clusters detected in lung lobes corresponding to the leukocyte subsets included alveolar, interstitial, and monocytic macrophages, neutrophils, dendritic cells (DCs), T and natural killer (NK) cells, and B cells. The results showed that in T492 virus infected hamsters, the peak of lung inflammation (Figure 6A) coincided with the influx of total leukocytes recruitment at 5 dpi (Figure 6I), and the cell counts of alveolar and monocytic macrophages in the T492 group were significantly higher than those in the 492I group (Figure 6J). This data indicate that the recruitment of inflammatory macrophages plays a critical role in lung damage during infection. Further, we tested the expression levels of monocytic macrophage-specific genes associated with viral RNA according to our previous report [49]. This gene set contains a range of chemokines, such as the CC subfamily and the CXC subfamily members. The results showed that 492I virus induced significantly lower levels of viral RNA associated chemokines in monocytic macrophages at 3 dpi when compared with control (Figure 6K). This reveals that a set of chemokines bursting at the early infection stage (3 dpi) is responsible for the recruitment of inflammatory macrophages at later disease stage (5 dpi). In this context, the attenuated viral pathogenicity of T492I mutation could be partly explained by the suppression of viral RNA associated chemokines in monocytic macrophages.

Next, we performed statistical analyses of sequenced strains with patient information (“Clinical_Info.xls” in Supplemental Data, from GISAID). The vaccinated cases were excluded from the analysis to avoid any potential distraction. We compared the ratios of pairs of opposite disease status (Table S5) between T492 and 492I variants collected in a six-month interval after the spread of Delta. The comparisons were in two geographic scales, country and region subdivision, to eliminate bias resulted from the difference of countries/regions. Despite the limited amount of data, we still found support for our experimental findings that 492I variant caused a lower ratio of severe illness than T492 variant in samples from two countries and two region-subdivisions (Figure 6L-O). Taken together, these data imply that the decreased disease severity of T492I may contribute to the attenuated pathogenicity of SARS-CoV-2 virus.

## DISCUSSION

Our study demonstrates the transmission advantage and adaptiveness of T492I by comprehensive in-silico analyses and experiments. The sub-variants of Delta (T492: 21A and 21I; 492I: 21J) provide nearly ideal samples for evolutionary analysis, because the influence of other mutations can be precluded. In this context, the evaluation with growth coalescent models are performed on the Delta variants with or without T492I. The NSP4 T492I substitution arises independently within multiple lineages (Figure 1A, B), suggesting a universal transmission advantage of this mutation among SARS-CoV-2 lineages. In the statistics of difference geographical levels, there are a few countries/regions showing a decreased IF of T492I in the epidemiogical survey, possibly due to the inadequacy of the sample size. Mechanistically, the T492I mutation is located at the interaction region between NSP4 and NSP5. We demonstrate that the T492I mutation increases the protease activity of NSP5 by enhancing the enzyme-substrate contact. The increased protease activity ultimately leads to the release of more NSPs cleaved by NSP5. For example, NSP6 contains ER-zippering activity and acts as an organizer of DMV clusters [31], NSP12 is a RNA-dependent RNA polymerase with nucleotidyltransferase activity, and NSP13 is a helicase with RNA 5’triphosphatase activity [21]. The elevated expression of these NSPs partly contributes to an increased amount of CMs and results in a strengthened replication and infectivity associated with the T492I mutation.

The IF of T492I, as well as the predicted relative fitness of the 492I variants over T492 variants are both increasing continuously. These results suggest the importance of T492I mutation to the rapid spread of 492I-bearing VOCs after the beginning of global vaccination program. SARS-CoV-2 evolved VOCs with immune evasion capability in the ongoing of the global vaccination program. Actually, the VOCs bearing 492I (Lambda, Mu, Delta and Omicron) all show immune evasion capability [50–53]. Vaccination may have accelerated the spread of T492I since vaccinated individuals could be susceptible to the strains with immune evasion capability, according to previous clinical data [54]. In our experimental data, NSP15 and NSP16, two NSPs cleaved by NSP5 that assist the viral immune evasion, had significantly higher protein levels in 492I group than in T492 group. Accordingly, SARS-CoV-2 variant bearing the T492I mutation showed a reinforced ability to evade antibody neutralization. Recently, NSP6 ΔSGF mutation has been reported to be associated with immune evasion capability [31]. Considering that all Omicron sub-variants bear both NSP4 T492I and NSP6 ΔSGF mutations, we hypothesize that the co-occurrence of these two mutations may contribute to a rapid increase in infectivity and elevated immune resistance in Omicron variants [51,55]. However, further experimental verification of this hypothesis is required.

Importantly, we found that hamsters infected with 492I virus exhibited less lung damage than those infected with T492 virus (Figure 6A, B), and the available clinical data support the conclusion gleaned from animal experiments (Figure 6L-O). The T492I mutation conferred an enhancement on viral infectivity, replication and immune resistance, but attenuated the pathogenicity. Similarly, mutation of S78R-K79R-S81R in Japanese Encephalitis Virus increased the cleavage of prM Protein, promoting viral replication but attenuating virulence [56]. These indicate that T492I may be the mutation that contributes to the attenuation of virulence in Omicron sub-variants. Considering that the spike mutation does not function on the virulence attenuation of Omicron [29] and nucleocapisid mutations have an enhancing effect on virulence [14], our study suggests that non-structural mutations in Omicron may contribute to the virulence attenuation predominantly. We hypothesize two possible mechanisms to explain the virulence attenuation. First of all, some of the NSPs cleaved by NSP5, such as NSP9, NSP10 and NSP11, may be involved in unknown physiological processes that reduce the virulence of virus; Secondly, it is possible that T492I mutation affect factors other than NSP5 activity, such as reducing the expression of factors known to be associated with virulence, including NSP1 and NSP3 [25–27], or affecting the binding of NSP4 to certain host proteins, thereby attenuating the pathogenicity. Nevertheless, the specific mechanisms underlying the changes of viral properties require further in-depth investigations. Our study here highlights the importance of NSP4 mutation in altering the properties of SARS-CoV-2 during the past three years of evolutionary process, and provides a potential strategy for screening antiviral drugs applicable to a variety of coronaviruses.

## ACKNOWLEDGMENTS

We gratefully acknowledge the submitting and the originating laboratories where genetic sequence data were generated and shared via NCBI and the GISAID Initiative. This work was supported by grants from the National Natural Science Foundation of China, SGC’s Rapid Response Funding for COVID-19 (C-0002), the National Natural Science Foundation of China (81970008, 32170661 and 82000020), the Fundamental Research Funds for the Central Universities (2020CDJYGRH-1005 and 2021CDJYGRH-009), Chongqing Talents: Exceptional Young Talents Project (No. cstc202lycjhbgzxm0099) and the Youth Innovative Talents Training Project of Chongqing (CY210102). The funders had no role in study design, data collection and analysis, decision to publish, or preparation of the manuscript.

## AUTHOR CONTRIBUTIONS

H.W., Z.Z., X.L. and C.J. collected the data and performed the population genetic analyses. H.W., X.L., Z.S., W.X., J.T., D.K. and Y.X. performed the experiments. W.X. and B.X. performed the protein structure analysis. H.W. conceived the idea. H.W., Z.Z., X.L. and Z.S. wrote the manuscript. H.W., W.X. and Z.Z. coordinated the project.

## MATERIALS AND METHODS

### Cell lines, animals and infection

Human lung adenocarcinoma epithelial Calu-3 cells (HTB-55, ATCC, MD, USA) and African green monkey kidney epithelial Vero E6 cells (CRL-1586, ATCC) were maintained at 37□°C with 5% CO_2_ in high-glucose Dulbecco’s modified Eagle’s medium (DMEM, Gibco, CA, USA) supplemented with 10% FBS (Gibco). Cells were infected with a multiplicity of infection (MOI) of 0.01 at the indicated time points. Six-week-old golden Syrian hamsters (Mesocricetus auratus) were obtained from Janvier Laboratories (Le Genest, St Isle, France). Wild type SARS-CoV-2 (USA_WA1/2020 SARS-CoV-2 sequence, GenBank accession No. MT020880) virus (T492) was generated by using a reverse genetic method as previously described [14,57,58]. NSP4 T492I mutation (492I) was introduced to wild type virus by overlap-extension PCR as previously described [14]. Hamsters were anaesthetized by isoflurane and intranasally infected with 2×10^4^ PFU virus. For the competition experiment, 12 hamsters received a 1:1 mixture of T492 and 492I virus. Hamsters were weighed and recorded daily. On day 3 and day 6 post infection, cohorts of 6 infected hamsters were anaesthetized with isoflurane, and nasal washes were collected with sterile PBS. The hamsters were then humanely euthanized immediately, and trachea and lung lobes were obtained as previously described [38]. This study was carried out in strict accordance with the Guidelines for the Care and Use of Animals of Chongqing University. All hamster operations were performed under isoflurane anaesthesia to minimize animal pain. The SARS-CoV-2 live virus infection experiments were performed under biosafety conditions in the BSL-3 facility at the Institut für Virologie, Freie Universität Berlin, Germany and performed in compliance with relevant institutional, national, and international guidelines for care and humane use of animal subjects.

### Plaque assay, viral subgenomic RNA assay and genomic RNA assay

Plaque assays were performed as previously described [14]. Briefly, approximately 1×10^6^ cells were seeded into 6-well plates, and cultured in 5% CO_2_ at 37□°C for 12 h. T492 or 492I virus was serially diluted in DMEM containing 2% FBS, and 200 μL aliquots were supplemented into cells. Cells were co-incubated with virus and then supplemented with overlay medium that contains *1%* SeaPlaque agarose (Lonza Bioscience, Basel, Switzerland). After 2 days of incubation, neutral red (Sigma-Aldrich, MO, USA) was used to stain the plates, and plaques were calculated. Viral subgenomic RNA assay was performed with primers that target envelope protein (E) gene sequences [14,59]. Viral genomic RNA assay was performed with primers that target Orf1ab sequences as previously described [14,60]. Briefly, total RNA of infectious cell lysate was extracted using an RNeasy Mini Kit (QIAGEN, Hilden, Germany). RT–PCR was performed using a Superscript III One-Step RT-PCR kit (Thermo Fisher Scientific, CA, USA) and an ABI StepOnePlus PCR system (Applied Biosystems, CA, USA) according to the manufacturer’s instructions.

### The inference of relative fitness based on changes in IF of variants

According to the theory of classical population genetics [61], for two genotypes A and B in a population, the change in proportion (Δp) is:

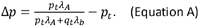

Here, *p_t_* and *q_t_* are the proportions of A and B in the population at the moment t, respectively. *λ_A_* and *λ_b_* are the absolute fitnesses of A and B, respectively. We manipulated the equation and obtained:

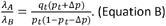

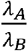 is the fitness of A relative to B at the moment t. Based on Equation B, we inferred the relative fitness of Delta 492I relative to Delta T492 variants. The time interval is a six-month interval from January 1^st^ 2021, the time point near the beginning of the global vaccination program (Figure S3A). We counted the weekly median of the relative fitness per day. We obtained the records of vaccination coverage (people vaccinated per hundred) in different regions from Our Word In Data (ourworldindata.org) and performed a correlation test between the weekly median of the relative fitness and the vaccination coverage in different geographic levels. We also built a maximum likelihood estimation line of the fitness trend and evaluated the trend by Mann-Kendall trend test. We performed binomial test for the significant cases in each geographic level.

### Function prediction based on clinical data

We manually gathered the clinical information of 115032 SARS-CoV-2 strains (Clinical_Info.xls in Data S1) from GISAID. Following the pipeline in our previous work [7], we grouped this information into pairs of opposite patient statuses, according to a series of keywords (Table S5). We counted the number of variants with different patient statuses and tested the significance by using the chi-squared test.

### Statistical Analysis

Sample size was based on empirical data from pilot experiments. The investigators were blinded during data collection and analysis. A value of P < 0.05 was considered significant.

## Supplemental Materials

Supplemental Material file includes:

Materials and Methods

References

Supplemental Figures S1-S3

Legends for Supplemental Tables S1-S5

---

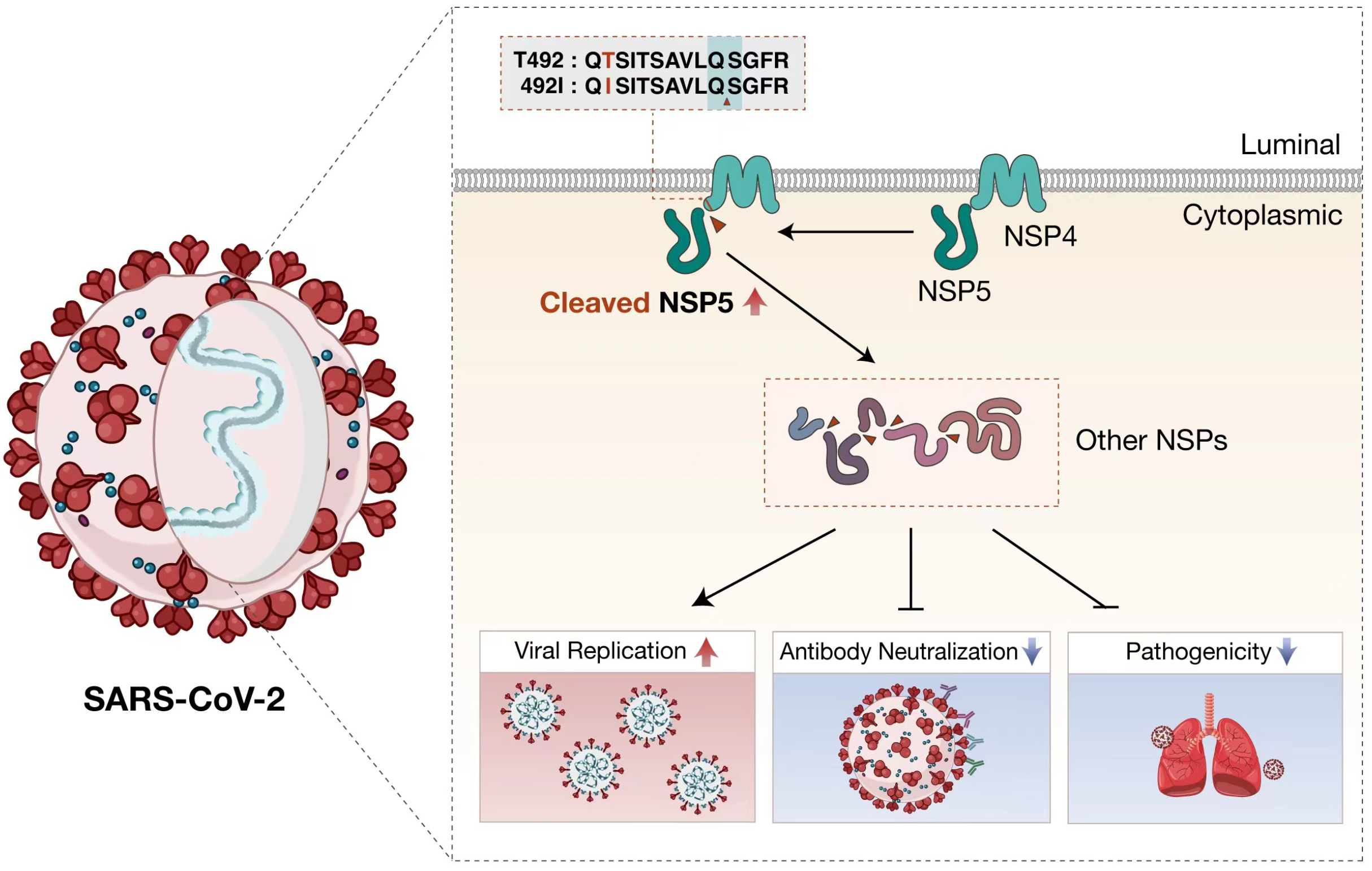

