## Supplemental Figure S1-S3 for "T492I mutation alters SARS-CoV-2 properties via modulating viral non-structural proteins"

**This file includes:**

Materials and Methods

References

Supplemental Figures S1-S3

Legends for Supplemental Tables S1-S5

### MATERIALS AND METHODS

#### Statistics of SARS-CoV-2 mutations and VOCs

We used the strain MN908947, collected in December 2019 [1,2], as the reference. We performed pairwise alignments of the protein sequences of the reference strain and all sequenced strains, downloaded from GISAID (www.gisaid.org, updated at September 22, 2022). The alignments are used to identify the genomic position of mutations. Based on the strain information of the phylogenetic tree provided by the Nextstrain database, we extracted the sample sequences of VOCs and built a library of the site substitutions of VOCs. The library was used to identify the attributes of all the sequenced strains. In the library, there are 288 substitution sites. We built shortened sequences containing the 288 sites using the alignments of all strains. The IF evaluation of the NSP4 T492I and VOCs was based on these shortened sequences. We flowed a reported approach [3] to compare the growth rates of different lineages. We wrote Perl scripts to count the weekly running counts of the strains with T492I and those without T492I in a six-month interval from January 2021 to June 2021. In the evaluation of the changes of 492I relative to T492 variants, we performed Fisher's exact test of the fraction of pairs of lineages on the onset, when there is an introduction of a new variant, and the day after more than two weeks. For a sufficient number of samples in statistics, we required that the sum of the counts of both lineages is higher than 20 at each time point. We built a maximum likelihood estimation line of the fraction trend and evaluated the trend by Mann-Kendall trend test and by isotonic regression analysis. These evaluations were performed in three hierarchical geographic levels (world, country and region subdivision). We used the same method to evaluate the IF of T492I in Delta variant in a six-month interval from April 2021 to September 2021. Mann-Kendall trend test was performed by the R package "trend". Wilcoxon test and Fisher's exact test were performed by R.

Following a reported pipeline [4], we evaluated the transmission advantages of 492I Delta variants compared to T492 variants in the California State, USA. A phylogenetic tree was constructed using all SARS-CoV-2 sequences collected in the California State from April 2021 to September 2021. The phylogenetic clusters were identified by TreeCluster [5] with the parameter -t 0.045. The simulations in a logistic growth coalescent model and in an exponential growth coalescent model were performed using an HYK substitution model, a strict clock type, an exponential prior distribution and a length of chains 10e7 by BEAST v1.10.4 [6]. We used FastTree [7] (parameters, -boot 1000) to build the phylogenetic trees of 492I Delta and T492 Delta variants, respectively. Then, we performed phylodynamic inference of effective population size for time-scaled phylogenies by the R package Skygrowth [8] (parameters, res = 300 and tau0 = 0.1). The plotting and analysis of the simulation data were performed by R libraries "gdata" and "ggplot2".

**Human airway tissue culture infection**

Viral infection in a primary human airway tissue culture was performed as previously described [9]. T492 or 492I virus was inoculated into a primary human airway tissue culture at a MOI of 5. From day 1 to day 5 after infection, sterile PBS was added on the apical side of the tissue culture and incubate for 30 min at 37°C to elute released virus.

**Neutralization assay**

Neutralization assay was performed using wild type and 492I viruses containing a mNeonGreen reporter as previously described [9,10]. Briefly, Calu-3 cells were plated in 96-well plates. The next day sera were serially diluted and incubated with mNeonGreen virus for 1 h. The virus-serum mixture was then transferred into a cell plate at a final MOI of 2. Infection rate was determined by dividing the number of mNeonGreen-positive cells by the total number of cells. The half-maximal neutralization titres ($NT_{50}$) was calculated using a nonlinear regression method. GraphPad Prism 8 (Graph Pad Software, CA, USA) was used to plot the curves of the relative infection rates versus the serum dilutions ($log_{10}$ values).

**Competition assay**

Competition assay was performed by RT-PCR with quantification of Sanger peak heights as previously described [9]. Briefly, a pair of common primers (Primers: SARS-CoV-2 9840F, 5'-AGTTGCGTAGTGATGTGC-3'; SARS-CoV-2 10296R, 5'-GAATGTCCAATAACCCTGA-3') was used to quantify the T492: 492I ratios. The 457 bp product was amplified using a SuperScript III One-Step RT–PCR kit (Thermo Fisher Scientific), followed by Sanger sequencing (Primer for Sanger sequencing: 5'-AGTTGCGTAGTGATGTGC-3'). The electropherograms were scored by QSVanalyser software to determine the T492: 492I ratios.

**Pathological examination and scoring**

Pathological examination of hamster tissues was performed according to standard procedures. Briefly, hamsters were anaesthetized with isoflurane and tissues were harvested. Tissues were fixed in 10% formalin, and trimmed and embedded in paraffin for hematoxylin and eosin staining. The histopathology scoring of the infected hamsters was performed using a semi-quantitative pathology scoring system as previously described [9]. Five sections were collected from each hamster, and the scores of lung sections were summed to calculate a total score of each animal. All scoring was performed by the same operator to ensure consistency of scoring.

**Pulmonary function test**

Pulmonary function test was performed on hamsters infected with T492 or 492I virus and control group (naïve, 0 dpi) following a previous report with some modifications [11,12]. Briefly, hamsters were anaesthetized with a mixture of sodium pentobarbital (3 mg/100 g), α-chloralose (3.8 mg/100 g) and urethane (38 mg/100 g). Then hamsters were tracheostomized and cannulated with a 16GA catheter. Measurements of functional residual capacity, total lung capacity, vital capacity, residual volume, forced expiratory volume in in 100 ms were performed using previously described settings[11,12].

**Protein structure modeling and docking of NSP5 and substrates**

Starting from the crystal structure of NSP4|5 peptidyl substrate (TSAVLQSGFR) (PDB ID: 7DVP) [13]. The 3D structures of the T492 (QTSITSAVLQSGFR) and 492I (QISITSAVLQSGFR) substrates cleaved by SARS-CoV-2 NSP5 in this work were constructed using RosettaRemodel method [14]. The blueprint file was first obtained from the crystal structure of nsp4|5 peptidyl substrate. The structure was then locally refined by inserting four residues (QTSI for the T492 and QISI for 492I) on N-terminal, still using loop fragments. The sidechains of this newly built loop segment were designed automatically and generated 100 conformations. According to the score file, the first conformation was selected for the following computing. The 3D structure of SARS-CoV-2 NSP5 (PDB ID: 7DVP) [13] was retrieved from PDB database. Before docking, the mutation (H41A) in the crystal structure was mutated back to its native state using the mutagenesis tool in PyMOL. NSP5-substrate docking was performed by RosettaDock [15]. The structures of the two substrates and NSP5 were firstly formed by the script of *clean_pdb.py*, and were refined by running relax protocol. Then, the structures of T492 and 492I substrates were manually docked to the relaxed structure of NSP5 using the nsp4|5 peptidyl substrate as a reference. The obtained model was further prepacked and used as a starting point for several rounds of local docking to generate 10,000 decoys by running docking with the Monte Carlo (MC) refinement algorithm. Finally, the energy funnel of each docking trajectory was depicted using the interface score (I_sc) and the interface root-mean-square deviation (I_rmsd).

**Immunoprecipitation and immunoblot assay**

Cell lysates were collected and lysed with RIPA Lysis Buffer (Merck Millipore, Darmstadt, Germany) containing protease inhibitor cocktail (Merck Millipore). Lysates were incubated with anti-His antibody (sc-8036, Santa Cruz Biotechnology, CA, USA) and Protein A-Agarose (Santa Cruz Biotechnology) overnight at 4°C. Precipitated protein complex was harvested and subjected to immunoblot assay. For immunoblot, samples were separated by sodium dodecyl sulfate polyacrylamide gel electrophoresis and transferred to polyvinylidene fluoride membranes (Merck Millipore). Blots were probed with anti-His (sc-8036, Santa Cruz Biotechnology), anti-Flag antibody (ab1162, Abcam, MA, USA) and visualized using a ChemiDoc imaging system (Bio-Rad, CA, USA).

**Dual luciferase reporter assay**

Dual luciferase reporter assay was performed using a dual luciferase reporter Assay System (Promega, WI, USA) according to the manufacturer’s instructions. HEK-293 cells were transfected with luciferase reporter plasmids, and internal control plasmid pRL-SV40 was cotransfected as an internal control. After 24 hours, cell lysates were harvested for dual luciferase detection using a VICTOR X5 Multilabel Plate Reader (PerkinElmer, Germany). Relative luciferase activity was measured by firefly luciferase luminescence divided by renilla luciferase luminescence.

**Enzyme linked immunosorbent assay**

Cell culture supernatants were purified by centrifugation, and then assayed by enzyme linked immunosorbent assay as previously described. Samples were probed with anti-NSP4 (SAB3501139, Merck Millipore), anti-NSP5 (SAB3501127), anti-NSP6 (SAB3501140), anti-NSP8 (SAB3501131), anti-NSP12 (NBP3-07963, Novus Biologicals, CO, USA), anti-NSP13 (NBP3-07055), anti-NSP15 (NBP3-11932), anti-NSP16 (A20283, ABclonal, MA, USA) antibodies. The concentration of each non-structural protein was calculated against a standard curve.

**Proximity ligation assay**

Proximity ligation assay was performed using a Duolink PLA Multicolor Probemaker kit (DUO96010, Sigma-Aldrich) according to the manufacturer's instructions. Briefly, Vero E6 cells were transfected with plasmids encoding Flag-tagged T492 or 492I substrates and then infected with SARS-CoV-2 virus. Cells were fixed with 4% PFA on slides and permeabilized with 0.2% Triton X-100. Oligo-conjugated anti-Flag and anti-NSP5 antibodies were diluted with Probemaker PLA Probe Diluent. After incubation, ligation, amplification and washing, the proximity of NSP5-substrates were determined by fluorescence intensity.

**Isothermal titration calorimetry and Surface plasmon resonance**

Isothermal titration calorimetry (ITC) and Surface plasmon resonance (SPR) were performed to determine the affinity of NSP5-substrates as previously described [9]. For ITC, purified proteins were transferred to buffer containing HEPES (20 mM, pH 7.5), NaCl (100 mM), and 2-mercaptoethanol (2 mM) by HiTrap desalting column (GE healthcare, GA, USA). Titrations were performed by using a Microcal PEAQ-ITC calorimeter (Malvern Panalytical, Malvern, UK) and data were analyzed using the PEAQ-ITC analysis software. For SPR, purified NSP5 protein (20 μg/mL) was coupled on a CM5 Chip (GE Healthcare, IL, USA) and different doses of T492 or 492I substrates were injected for dissociation analysis. Dissociation rate constants were determined using the steady state affinities obtained for each enzyme concentration.

**Intracellular fluorescence resonance energy transfer**

Schematic diagram of intracellular fluorescent resonance energy transfer assay was shown in Figure S3 following a previous report with modifications [16]. HEK-293T cells were transfected with CMV-3×492I(TAG)-NSP5-YFP or CMV-3×T492(TAG)-NSP5-YFP plasmid, and then treated with 50 μM ANAP. After 18 hours, cells were transfected with different dose (0.5, 1.0 and 2.5 μg, respectively) of 3×T492 or 3×492I plasmid. A fluorescent amino acid ANAP was incorporated into the T492 or 492I substrate. The fluorescence of ANAP-substrate was detected with excitation at 405 nm, and the fluorescence of YFP-NSP5 was detected with excitation at 488 nm.

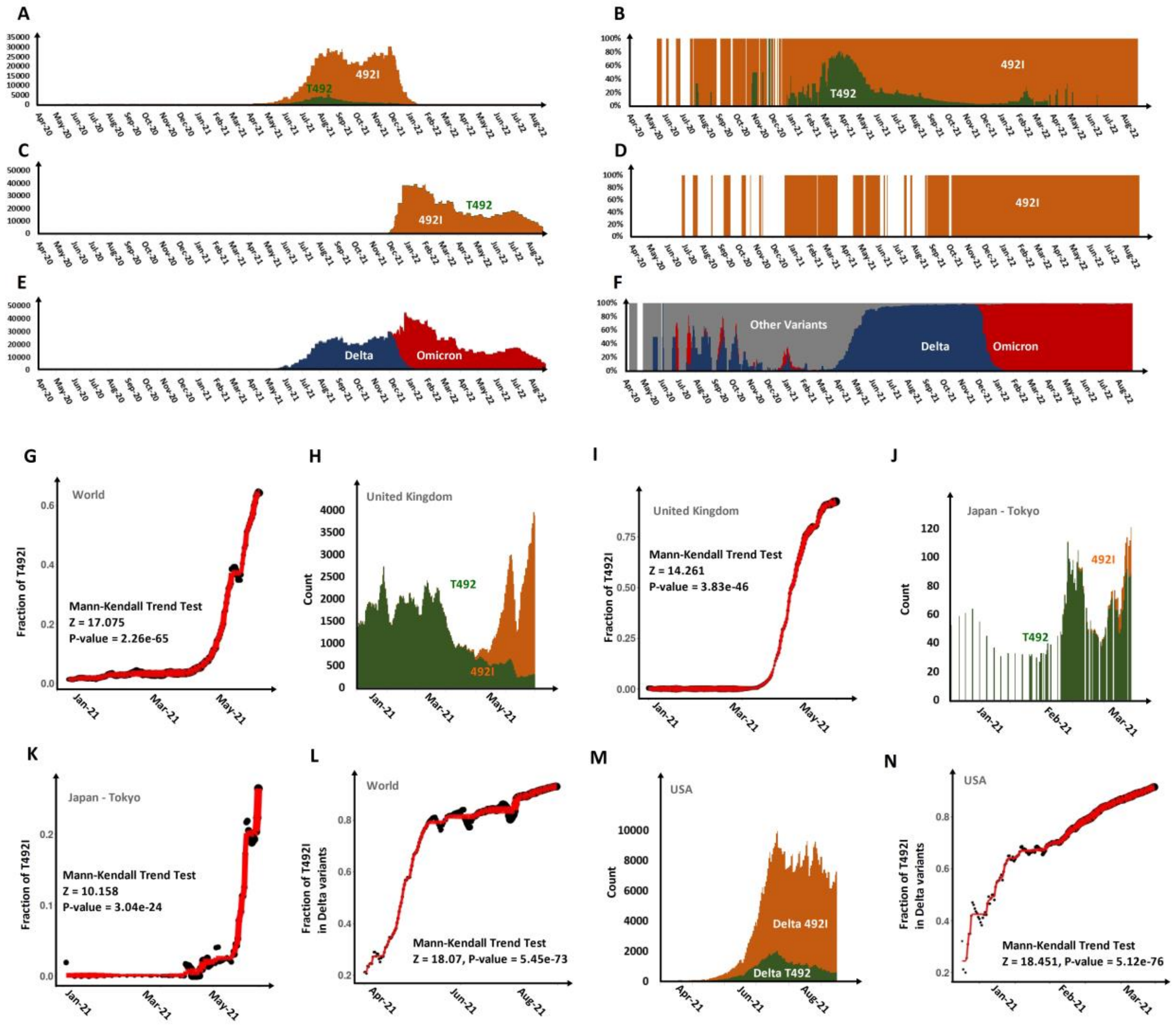


**Figure S1**. Additional evidences and samples supporting the adaptiveness of the NSP4 mutation T492I.

(A) The weekly running counts of T492 Delta variants and 492I Delta variants around the world from 2020 to date. (B) The changes of the fraction of T492I in Delta variants around the world from 2020 to date. (C) The weekly running counts of T492 Omicron variants and 492I Omicron variants around the world from 2020 to date. (D) The changes of the fraction of T492I in Omicron variants around the world from 2020 to date. (E) The weekly running counts of Delta and Omicron variants in T492I strains around the world from 2020 to date. (F) The changes of the fraction of Delta and Omicron variants in T492I strains around the world from 2020 to date. (G), (I) and (K) are the fitted trend of the change in the fractions of T492I in the world, United Kingdom and Japan-Tokyo, respectively. (L) and (N) are the fitted trend of the change in the fractions of T492I in Delta variants the world and USA, respectively. The legends in (G), (I), (K), (L) and (N) follow Figure 1H. (H) and (J) are the weekly running counts of T492 variants (dark green) and 492I variants (orange) from January 2021 to June 2021 and in United Kingdom and Japan-Tokyo, respectively. (M)The weekly running counts of Delta T492 variants (dark green) and Delta 492I variants (orange) from April 2021 to September 2021 in the USA.

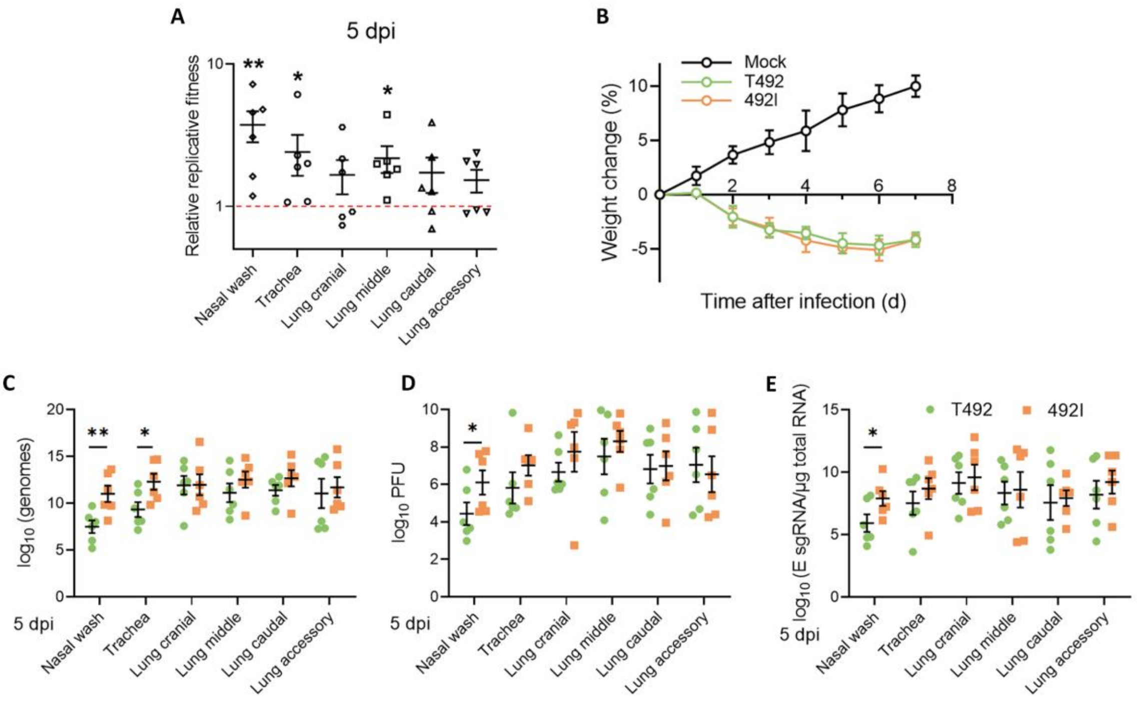


**Figure S2**. 492I virus has higher competitiveness and infectivity in hamsters. (A) Hamsters were infected with T492 and 492I virus at a mixture of 1:1. The relative amounts of T492 and 492I viral RNA in nasal wash, trachea and lung samples were detected by RT-PCR and Sanger sequencing at 5 dpi. Log10 scale was used for the Y-axis. Dots represent individual hamsters (n = 6). (B-E) Hamsters were infected with $2\times10^4$ PFU of T492 or 492I virus. Weight loss (B) was monitored for 7 consecutive days. n = 12 (all cohorts) at days 0–3; n = 6 (all cohorts) at days 4–7. Weight loss was analyzed by ANOVA with Tukey's post hoc test. Genomic RNA levels (C), PFU titres (D), and E sgRNA loads (E) in nasal wash, trachea and lung samples were detected at 5 dpi. Dots represent individual hamsters (n = 6). Data are presented as the mean ± s.e.m.. *, $p<0.05$, **, $p<0.01$.

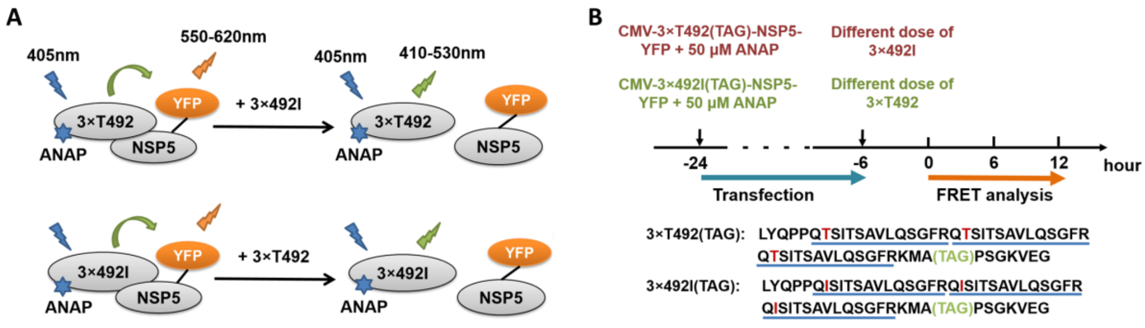


**Figure S3**. Schematic diagram of FRET analysis. (A, B) HEK-293T cells were transfected with CMV-3×492I(TAG)-NSP5-YFP or CMV-3×T492(TAG)-NSP5-YFP plasmid, and then treated with 50 μM ANAP. After 18 hours, cells were transfected with different doses (0.5, 1.0 and 2.5 μg, respectively) of 3×T492 or 3×492I plasmid. Graph displays the FRET ratios (IYFP/IANAP) recorded from single cell images. If the T492-NSP5 (or 492I-NSP5) complex is disrupted by competition, IYFP/IANAP ratio will decrease.

**Legends for Supplemental Tables S1-S5**

Table S1. Evidences supporting the transmission advantage of T492I in different geographic scales. The columns on the left are comparison of the fraction between T492 and 492I for two time points separated by a more than 2-week gap in different geographical scales. The first time point is the onset day. The second time point is more than 2 weeks after the onset date. Changes in fraction were evaluated by Fisher's exact test. The columns on the right are comparison of the growth rates by Mann-Kendall trend test (MK) and isotonic regression (IR). A positive MK Z-value with P-value < 0.05 indicates an increase in the fraction of Lineage 2 with a statistical significance. For IR analysis, the P-value was calculated for two one-sided tests by comparing the null hypothesis of no consistent changes in relative frequency over time with positive or negative pressure. Only the results with significance (P-value<0.05) were presented. At the country and region scales, Binomial P-value were provided against the null hypothesis that increases and decreases were equally likely.

Table S2. Countries/regions with evidence to support the transmission advantage of T492I in Delta variants. Legends Follow Table S1.

Table S3. Catalytic efficiencies for T492I substrates.

Table S4. Evalutaion of the trends and the correlation with the vaccination coverage (VC) for the fitness of 492I (Delta) relative to T492I Delta in different countries. Only the cases with significant coorelation or with significant MK test Z-value were listed. At the country and region scales, Binomial P-value were provided against the null hypothesis that increases and decreases were equally likely.

Table S5. Keywords used to search and annotate records with patient status.
